# Evaluation of exosome-encapsulated miR-23a/b in the diagnosis of human coronary heart diseases

**DOI:** 10.1101/2020.12.27.424501

**Authors:** Changzhi Xu, Hui Xiao, Yanhua Yi, Donglin Zhu, Xiaojing Yue, Yun Xi

## Abstract

**Objective:** Coronary heart disease (CHD) is currently one of the major causes of death with high morbidity. Due to the increasing heterogeneity and complexity in the CHDs progression, biomarkers for specific diagnosis and monitoring of disease progression need to be developed. The study was aimed to investigate the roles of serum exosomal miR-23a and miR-23b in diagnosis of CHDs.

**Methods:** 16 healthy individuals and 56 patients with CHDs were enrolled in this study, including the CHDs of stable angina, unstable angina, non-ST elevation myocardial infarction (NSTEMI), ST elevation myocardial infarction (STEMI) and acute myocardial infarction (AMI). Serum exosomal miR-23a and miR-23b were quantified by Q-PCR. The associations of miR-23a/b with multiple clinical parameters were analyzed.

**Results:** Serum exosomal miR-23a was downregulated in the 56 CHD patients. In the specific prediction of stable angina and AMI, miR-23a achieved the area under the receiver operating characteristic (AUC-ROC) of 0.809 and 0.783, respectively. The levels of serum creatinine (CREA) and miR-23a were associated, which was consistent with risk of kidney injury in CHDs patients. Exosomal miR-23b levels showed no difference among CHD groups.

**Conclusions:** miR-23a may serve as a non-invasive marker in predicting stable angina and AMI. The level of miR-23a was implicated in CREA-dependent CHD-renal damage.

## Introduction

Coronary heart diseases (CHDs), including stable angina, unstable angina, myocardial infarction and sudden cardiac death, affect 2-6% of the general population[1]. According to the data from National Health and Nutrition Examination Survey (NHANES), in 2015, CHDs mortality was 366801 in the US[2]. Although the treatments for CHDs were greatly improved in the past decades, the 5-year survival rate for CHDs still has not made progress. Due to the increasing heterogeneity and complexity in the CHDs progression, biomarkers for specific diagnosis and monitoring of disease progression need to be developed.

Exosomes are small extracellular vesicles with 50-90 nm in diameter, released by the fusion of intracellular multivesicular bodies (MVB) with the plasma membrane and diffused into the extracellular medium. In these processes, exosomes are capable of carrying signaling molecules including mRNA, miRNA and protein, and can mediate the complex exchange between cells[3]. The molecules that can be packaged within exosomes appear to reflect the diversity and signaling potential of these vesicles[4]. Numerous studies investigated the roles of exosomes in many different areas such as cancer and cardiovascular diseases, and proved that the contents of exosomes potentially be reflective of disease state[5; 6; 7]. Exosomes have been proved to be detectable in nearly all body fluids including blood[8], urine[9], ascites[10], cerebrospinal fluid[11] and saliva[12], and thus might be served as potential candidates for non-invasive biomarkers that indicative of real-time disease state. In this study, exosomes were isolated from the serum of CHDs patients, and the potential roles of exosome-encapsulated miR-23a and miR-23b in CHDs diagnosis were investigated.

microRNAs are short non-coding RNAs of 21-23 nucleotides playing important roles in various biological processes. In previous studies both miR-23a and miR-23b (miR-23a/b) were critical miRNAs associated with carcinogenesis[13; 14]. miR-23a might serve as a biomarker in the diagnosis and prognosis of multiple human cancers [15]. The expression pattern of miR-23a has been proved to be associated with tumor infiltration, tumor differentiation degree and lymph node metastasis[15]. Based on cellular experiments, the recent studies revealed that miR-23a/b participated in the hypoxic condition and chemicals induced cardiomyocyte apoptosis and growth, which suggested its roles in cardiac ischemic-reperfusion injury and myocardial hypertrophy[16; 17; 18; 19; 20; 21; 22]. However, the association of miR-23a/b and cardiovascular diseases was very limited studied. In this study, we assessed exosomal miR-23a/b expression pattern in the serum of 16 healthy individuals and 56 patients with CHDs. We revealed that the downregulated miR-23a was significantly associated with CHDs. After further analysis of the CHDs subjects based on the clinical characteristics, we demonstrated that miR-23a emerged as a potential non-invasive biomarker for the specific diagnosis of stable angina and acute myocardial infarction (AMI), respectively.

## Methods

### Patient recruitment

The study was approved by the Ethical Committee of the institute and all subjects were obtained with written informed consent. Participants were 16 healthy volunteers (Group 1), 16 patients with CHD-stable angina (Group 2), 8 patients with CHD-unstable angina (Group 3), 8 patients with ST elevation myocardial infarction (STEMI, Group 4), 8 patients with non-ST elevation myocardial infarction (NSTEMI, Group 5), and 16 patients with acute myocardial infarction (AMI, Group 6). Healthy volunteers were all non-smokers and had no history of chronic alcohol consumption, hypertension, diabetes or other medical disease.

### Serum Preparation and exosome isolation

After centrifugation of peripheral blood at 2000 x g for 15 minutes, the serum in supernatant was collected in a RNase-DNase free screw-cap vial. Total exosomes were extracted using EVast Exosome Precipitation Solution (EVast-S, Biotron) according to the manufacturer’s protocol. Briefly, 240μl serum was processed by centrifugation for 15 minutes at 3000 x g at 4℃ to remove cell debris. The supernatant was mixed with 60μl EVast Exosome Precipitation Solution, and the mixture solution was kept standing on ice for 30 minutes followed by being centrifuged at 1500 x g for 30 minutes at 4℃. The exosome-containing pellet was washed once with 500μl phosphate buffered saline (PBS) and re-suspended in 50μl PBS for immediate use.

### Reverse transcription and microRNA assays

Total exosomal RNA was prepared by TRIzol extraction (TRIzol Reagent, 15596026, Life Technologies). 20pmol cel-miRNA-39 and microRNA-negative control (miRNA-NC, UUGUACUACACAAAAGUACUG) served as controls, respectively. The reverse transcriptions of microRNAs were performed using TaqMan™ MicroRNA Reverse Transcriptase Kit (4366596, Invitrogen). The miRNA-specific primers for reverse transcription and the TaqMan probes for real-time PCR were from Taqman microRNA assays (4440886, hsa-miR-23b-3p, Assay ID 245306_mat; 4427975, hsa-mir-23a-3p, Assay ID 000399; 4427975, cel-miR-39-3p, Assay ID 000200, Thermofisher). The relative expression of individual miRNA was calculated as ΔCt = mean Ct miRNA-mean Ct miR-39.

### Statistical analysis

Baseline characteristics were presented as mean and standard deviation (SD) for continuous variables, or proportions for categorical variables. Patient characteristics were compared using the Student’s *t*-test. Mann-Whitney U test was used in the comparisons of miRNA level between control and CHD group. Pearson Correlation Coefficient was used to analyze correlations between miRNA level and clinical parameters. Receiver operating characteristic (ROC) curve was used to analyze the accuracy of miR-23a of predicting stable angina and AMI. A *P*-value of < 0.05 was considered significant. All statistical analyses were performed using SPSS 22.0.

## Result

### Clinical characteristics

Overall, 16 healthy controls and 56 CHDs patients including stable angina, unstable angina, STEMI, NSTEMI and AMI were recruited. The levels of exosomal miR-23a and miR-23b were detected in the subjects from control and 5 different CHD groups. Each group included 8 individuals and the baseline characteristics were summarized in Table 1. The fasting blood glucose was significantly increased in the patients with NSTEMI (8.19±2.65 vs 5.42±0.67 mmol/L, *P* = 0.0124) or AMI (6.78±1.36 vs 5.42±0.67 mmol/L, *P* = 0.0237), suggesting the potential contributor of diabetes. HDLC was significantly downregulated in the patients with stable angina (1.01±0.25 vs 1.36±0.29 mmol/L, *P* = 0.0216), unstable angina (0.98±0.17 vs 1.36±0.29 mmol/L, *P* = 0.0065) and NSTEMI (0.88±0.19 vs 1.36±0.29 mmol/L, *P* = 0.0016), indicating the role of cholesterol in the pathogenesis of CHDs. The NSTEMI patients displayed higher levels of blood urea nitrogen (8.24±3.2 vs 5.21±1.26 mmol/L, *P* = 0.0259) and serum uric acid (UA) (505±129 vs 356±138 μmol/L, *P* = 0.0426), suggesting that the kidney injury was likely to be more severe in the NSTEMI group.

**Table 1.** Comparison of clinical characteristics between healthy controls and CHDs patients. STEMI indicates ST elevation myocardial infarction; NSTEMI, non-ST elevation myocardial infarction; AMI, acute myocardial infarction; HDLC, high density liptein cholesterol; LDLC, low density liptein cholesterol; APoA, Apolipoprotein A; APoB, Apolipoprotein B; LPa, Serum lipoprotein a; and Cys C, cystatin C; \**P*<0.05.

|  | Group 1<br>Healthy<br>control | Group 2<br>CHD-stable<br>angina | Group 3<br>CHD-unstable<br>angina | Group 4<br>STEMI | Group 5<br>NSTEMI | Group 6<br>AMI |
| --- | --- | --- | --- | --- | --- | --- |
| Number of subjects | 8 | 8 | 8 | 8 | 8 | 8 |
| Age (years),mean (SD) | 50 (4) | 63 (5) | 65 (14) | 57 (11) | 65 (12) | 61 (11) |
| Male, % | 50 | 50 | 75 | 87.5 | 100 | 87.5 |
| Female, % | 50 | 50 | 25 | 12.5 | 0 | 12.5 |
| Hypertension, % | 0 | 50 | 62.5 | 75 | 62.5 | 37.5 |
| Diabetes, % | 0 | 12.5 | 37.5 | 12.5 | 12.5 | 12.5 |
| Smoker, % | 0 | 38 | 37.5 | 50 | 25 | 62.5 |
| chronic alcohol consumption, % | 0 | 25 | 0 | 12.5 | 0 | 37.5 |
| Fasting blood glucose (mmol/L), mean (SD) | 5.42 (0.67) | 5.25 (0.90) | 6.06 (1.11) | 5.93 (1.49) | 8.19 (2.65) * | 6.78 (1.36) * |
| Serum total cholesterol (mmol/L), mean (SD) | 5.40 (0.69) | 4.44 (0.82) * | 4.47 (1.79) | 4.84 (1.38) | 4.73 (1.38) | 5.39 (0.95) |
| Triglyceride (mmol/L), mean (SD) | 1.64 (1.10) | 1.48 (0.59) | 1.55 (0.48) | 1.39 (0.87) | 2.49 (1.12) | 1.23 (0.30) |
| HDLc (mmol/L), mean (SD) | 1.36 (0.29) | 1.01 (0.25) * | 0.98 (0.17) * | 1.14 (0.61) | 0.88 (0.19) * | 1.06 (0.40) |
| LDLc (mmol/L), mean (SD) | 3.36 (0.77) | 2.90 (0.75) | 3.02 (1.53) | 3.30 (1.01) | 2.98 (1.01) | 3.69 (0.90) |
| APoA (g/L), mean (SD) | NA | 1.06 (0.20) | 1.25 (0.22) | 1.07 (0.38) | 1.08 (0.28) | 0.90 (0.30) |
| APoB (g/L), mean (SD) | NA | 1.00 (0.21) | 0.96 (0.35) | 1.01 (0.14) | 1.09 (0.33) | 1.19 (0.21) |
| LPa (mg/L), mean (SD) | NA | 352 (387) | 226 (233) | 205 (115) | 260 (461) | 398 (373) |
| Cys C | NA | 1.20 (0.25) | 1.20 (0.43) | 1.02 (0.41) | 1.71 (1.03) | 1.45 (0.76) |
| Serum creatinine (μmol/L), mean (SD) | 77.86 (18.24) | 94.82 (17.83) | 94.75 (33.32) | 89.75 (29.76) | 126.00 (67.15) | 110.50 (43.32) |
| Blood urea nitrogen (mmol/L), mean (SD) | 5.21 (1.26) | 5.90 (1.69) | 6.37 (2.64) | 5.47 (1.66) | 8.24 (3.20) * | 6.58 (3.63) |
| Serum uric acid (μmol/L), mean (SD) | 356 (138) | 474 (106) | 497 (174) | 363 (179) | 505 (129) * | 412 (91) |

The associations of miR-23a, stable angina and AMI were analyzed in the subjects of 16 healthy individuals, 16 stable angina patients and 16 AMI patients. The baseline characteristics were shown in Table 2. Similar as the data presented in table 1, HDLC was also downregulated in the stable angina patients (1.03±0.25 vs 1.44±0.34 mmol/L, *P* = 0.0005) and AMI patients (0.95±0.31 vs 1.44±0.34 mmol/L, *P* = 0.0002). Both elevated levels of serum creatinine (CREA) and serum uric acid were detected in the stable angina patients (90.7±19.77 vs 75.93±15.89 mmol/L, *P* = 0.0268; 437±106 vs 352±109 μmol/L, *P* = 0.0329) and AMI patients (113.81±48.49 vs 75.93±15.89 mmol/L, *P* = 0.0058; 501±211 vs 352±109 μmol/L, *P* = 0.0177), suggesting the kidney injury in these two CHD groups.

**Table 2.** Comparison of clinical characteristics between healthy control, CHD-stable angina and AMI groups. STEMI indicates ST elevation myocardial infarction; NSTEMI, non-ST elevation myocardial infarction; AMI, acute myocardial infarction; HDLC, high density liptein cholesterol; LDLC, low density liptein cholesterol; APoA, Apolipoprotein A; APoB, Apolipoprotein B; LPa, Serum lipoprotein a; and Cys C, cystatin C; \**P*<0.05.

|  | Group 1<br>Healthy<br>control | Group 2<br>CHD-stable<br>angina | Group 6<br>AMI |
| --- | --- | --- | --- |
| Number of subjects | 16 | 16 | 16 |
| Age (years),mean (SD) | 54 (6) | 62 (8) | 59 (13) |
| Male, % | 56.3 | 66.7 | 87.5 |
| Female, % | 43.7 | 33.3 | 12.5 |
| Hypertension, % | 0 | 33.3 | 31.3 |
| Diabetes, % | 0 | 16.7 | 18.8 |
| Smoker, % | 0 | 33.3 | 62.5 |
| chronic alcohol consumption, % | 0 | 22.2 | 18.8 |
| Fasting blood glucose (mmol/L), mean (SD) | 5.35 (0.60) | 5.42 (0.80) | 8.65 (7.96) |
| Serum total cholesterol (mmol/L), mean (SD) | 5.79 (0.96) | 4.27 (1.12) * | 4.99 (1.22) |
| Triglyceride (mmol/L), mean (SD) | 1.68 (0.89) | 1.77 (1.23) | 1.54 (0.57) |
| HDLc (mmol/L), mean (SD) | 1.44 (0.34) | 1.03 (0.25) * | 0.95 (0.31) * |
| LDLc (mmol/L), mean (SD) | 3.58 (1.09) | 2.55 (1.04) * | 3.48 (1.10) |
| APoA (g/L), mean (SD) | NA | 1.12 (0.19) | 0.97 (0.25) |
| APoB (g/L), mean (SD) | NA | 1.00 (0.30) | 1.10 (0.29) |
| LPa (mg/L), mean (SD) | NA | 335 (381) | 334 (279) |
| Cys C | NA | 1.39 (1.41) | 1.47 (1.05) |
| Serum creatinine (μmol/L), mean (SD) | 75.93 (15.89) | 90.70 (19.77) * | 113.81 (48.49) * |
| Blood urea nitrogen (mmol/L), mean (SD) | 5.29 (1.22) | 5.28 (1.45) | 7.41 (5.48) |
| Serum uric acid (μmol/L), mean (SD) | 352 (109) | 437 (106) * | 501 (211) * |

### Exosome-encapsulated miR-23a was detected in various CHDs groups

To investigate the specific role of miR-23a/b in different characterized CHDs, the CHDs subjects were summarized as 5 groups: stable angina, unstable angina, STEMI, NSTEMI and AMI. Each group contained 8 individuals. The level of miR-23a was detected in each group by real-time quantitative PCR analysis. The ΔCt values were obtained after the comparison with exogenous control cel-miR-39 and miRNA-NC. The ΔCt values of miR-23a were elevated in the patients with stable angina or AMI, when compared with the healthy controls (Figure 1A and Figure 1E), indicating that miR-23a was downregulated in these two groups. No differences were detected in the comparisons between control and unstable angina, USTEMI and STEMI groups (Figure 1).

**Figure 1.**
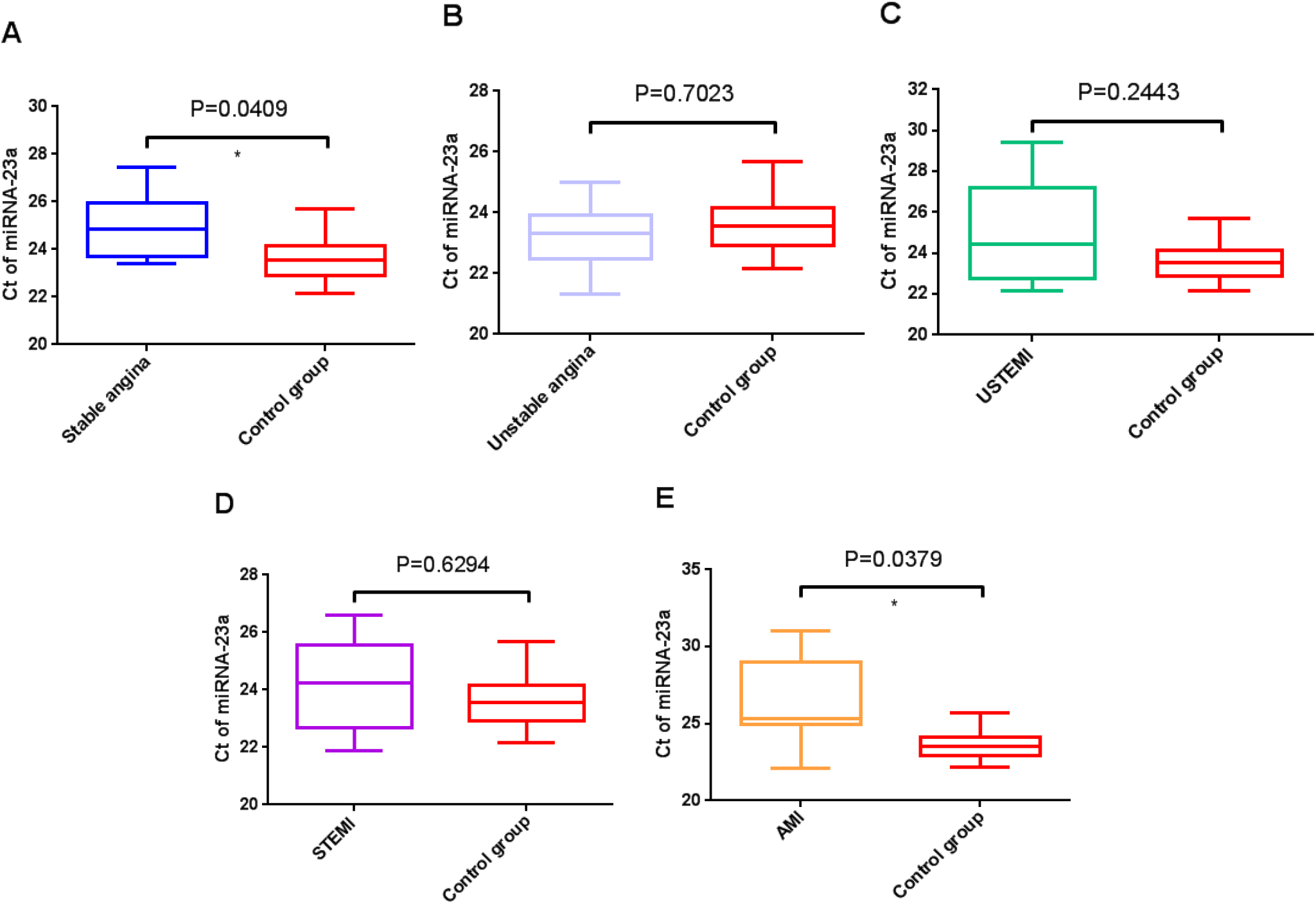
Box plots of exosomal miR-23a expression in healthy subjects and in differently classified CHDs patients. miR-23a was significantly downregulated in the serum exosomes from the patients with stable angina (**A**) and AMI (**E**). The levels of miR-23a showed no difference in unstable angina group (**B**), USTEMI group (**C**) and STEMI group (**D**). Each group included 8 individuals. STEMI indicates ST elevation myocardial infarction; NSTEMI, non-ST elevation myocardial infarction; AMI, acute myocardial infarction. \**P*<0.05

### Exosome-encapsulated miR-23b was detected in various CHDs groups

The serum exosomal miR-23b levels were detected in the control and 5 CHD groups via real-time quantitative PCR analysis. Among the 8 healthy control and 40 CHDs patients, no significant difference was found in the CHDs groups comparing to the control (Figure 2).

**Figure 2.**
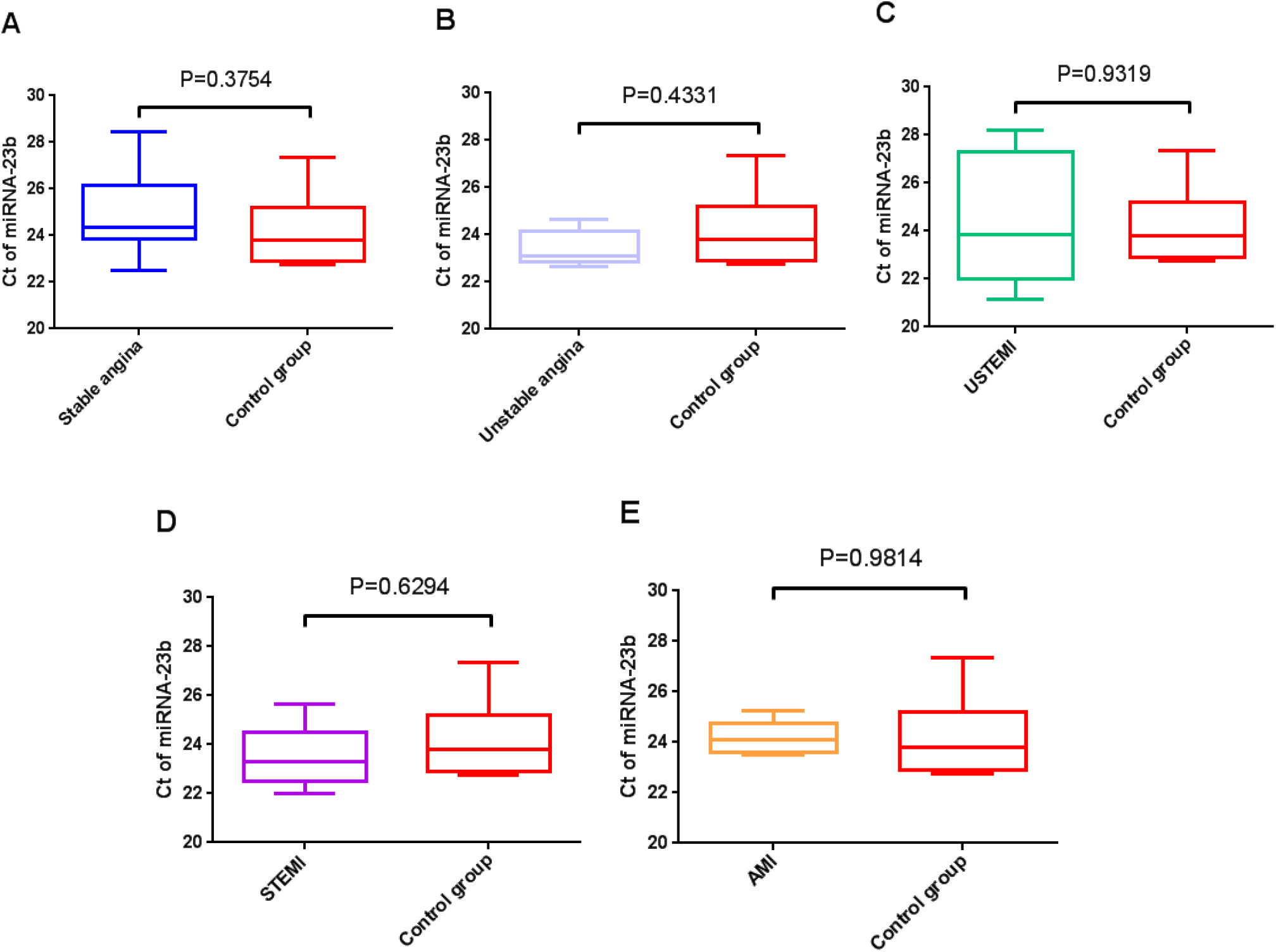
Box plots of exosomal miR-23b expression in healthy subjects and in differently classified CHDs patients. The expression of miR-23b in CHDs patients displayed no difference compared to the healthy control. Each group included 8 individuals. STEMI indicates ST elevation myocardial infarction; NSTEMI, non-ST elevation myocardial infarction; AMI, acute myocardial infarction.

### Serum exosomal miR-23a was downregulated in the patients with stable angina or AMI

To further explore the association of miR-23a with stable angina and AMI, more subjects were enrolled and each group contained 16 individuals. The clinical characteristics of 16 healthy control and 32 CHDs patients were presented in Table 2. The significant downregulation of serum exosomal miR-23a was confirmed in16 stable angina patients and 16 AMI patients, respectively (Figure 3). Based on the data of the stable angina and AMI groups, the correlations between serum exosomal miR-23a level and risk factors were analyzed, including gender, age, smoker, alcohol intake, diabetes and hypertension. No significant difference was found in these comparisons (Table 3).

**Figure 3.**
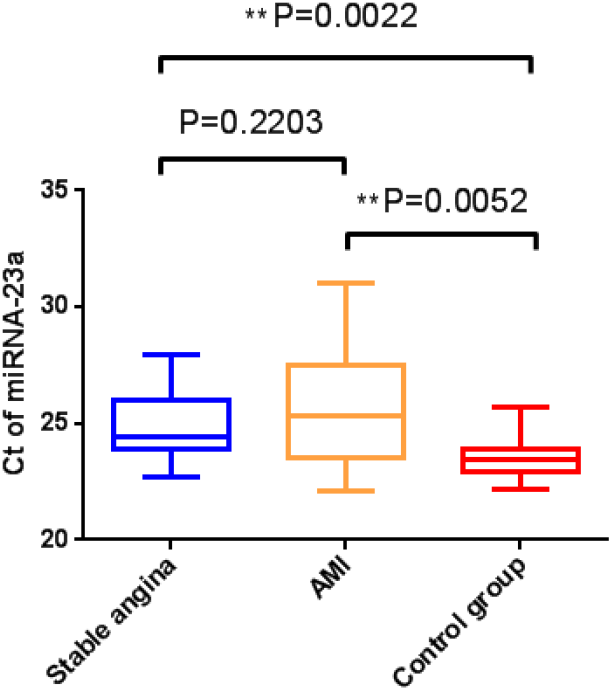
The downregulation of serum exosomal miR-23a expression was confirmed in the patients with stable angina or with AMI. Each group included 16 individuals. AMI, acute myocardial infarction. \*\**P*<0.05

**Table 3.** Correlation analysis between serum exosomal miR-23a levels and various clinical parameters of patients with stable angina or AMI. AMI indicates acute myocardial infarction.

| Characteristics | CHD-stable angina (N=16) |  |  | AMI (N=16) |  |  |
| --- | --- | --- | --- | --- | --- | --- |
|  | N | Ct of miR-23a<br>(mean±SD) | *P* value | N | Ct of miR-23a<br>(mean±SD) | *P* value |
| Gender |  |  |  |  |  |  |
| Man | 10 | 24.58±1.41 | >0.05 | 14 | 25.74±2.66 | >0.05 |
| Woman | 6 | 25.30±1.61 |  | 2 | 26.38±2.05 |  |
| Age (yr) |  |  |  |  |  |  |
| ≤55 | 3 | 23.99±1.13 | >0.05 | 5 | 24.79±1.71 | >0.05 |
| >55 | 13 | 25.04±1.51 |  | 11 | 26.29±2.78 |  |
| Smoker |  |  |  |  |  |  |
| No | 9 | 25.38±1.80 | >0.05 | 6 | 24.86±2.37 | >0.05 |
| Yes | 7 | 24.16±0.41 |  | 10 | 26.39±2.71 |  |
| Alcohol intake |  |  |  |  |  |  |
| No | 12 | 24.83±1.63 | >0.05 | 12 | 25.64±2.69 | >0.05 |
| Yes | 4 | 24.89±1.10 |  | 3 | 26.83±2.56 |  |
| Diabetes |  |  |  |  |  |  |
| No | 12 | 24.72±1.42 | >0.05 | 11 | 25.31±2.47 | >0.05 |
| Yes | 4 | 25.23±1.82 |  | 4 | 27.45±2.70 |  |
| Hypertension |  |  |  |  |  |  |
| No | 8 | 24.97±1.37 | >0.05 | 7 | 25.37±2.85 | >0.05 |
| Yes | 8 | 24.73±1.66 |  | 8 | 26.32±2.51 |  |

### Role of serum exosomal miR-23a as a predictor of stable angina and AMI

ROC curve was introduced to analyze the accuracy of the potential biomarker miR-23a on predicting stable angina and AMI. It was found that the area under the ROC curve (AUC) for predicting stable angina was 0.809 (95% CI = 0.654-0.963, *P* = 0.003), and the AUC for predicting AMI was 0.783 (95% CI = 0.606-0.960, *P* = 0.006) (Figure 4). These data indicated that miR-23a had the potential to be a promising biomarker in predicting the AMI and stable angina.

**Figure 4.**
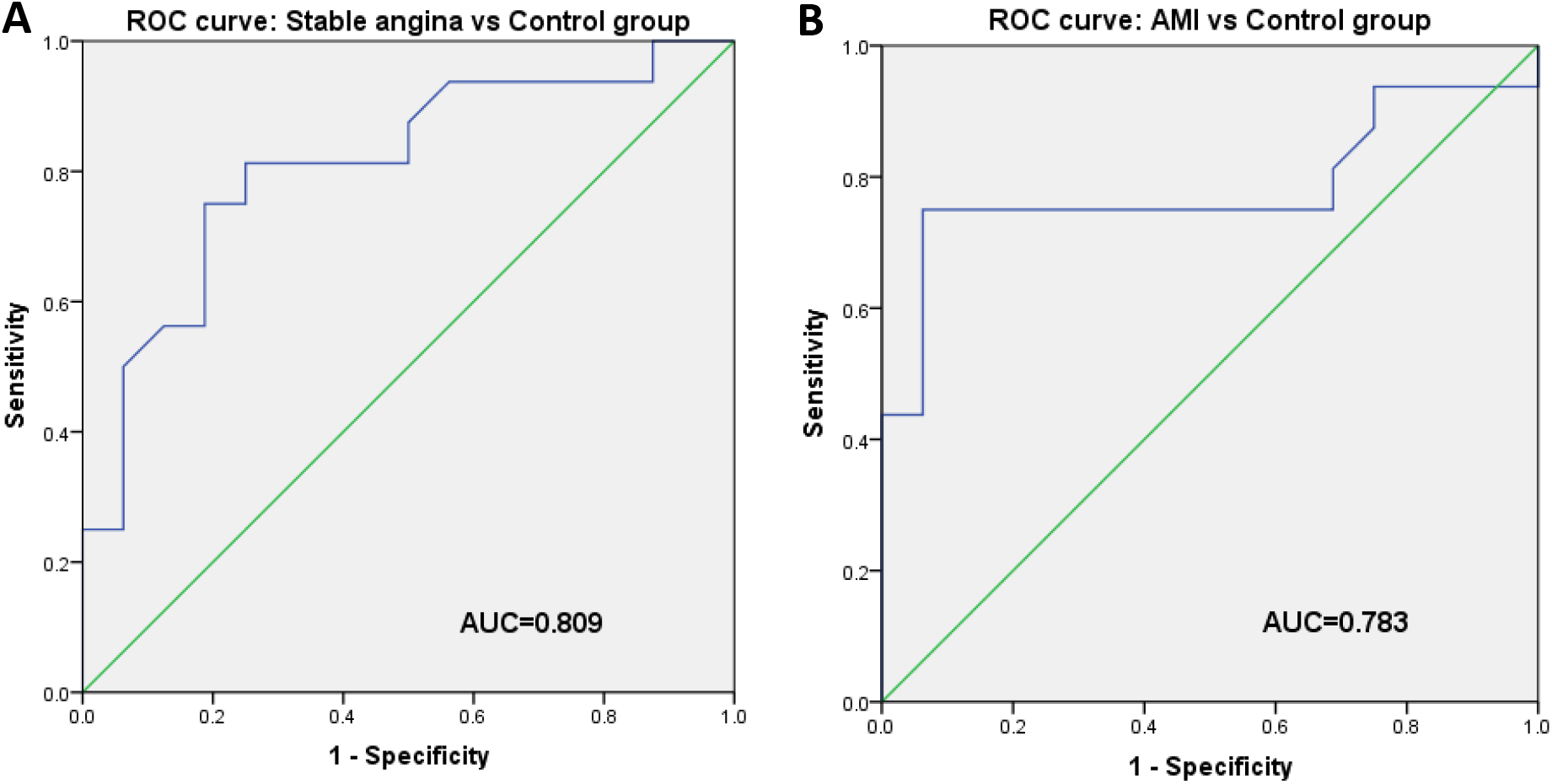
ROC curve of miR-23a on predicting stable angina and AMI. ROC indicates receiver operating characteristic; AMI, acute myocardial infarction.

### Correlation between serum exosomal miR-23a level and clinical parameters

Given that the clinical parameters HDLC, CREA and UA were significantly changed in the AMI and stable angina groups compared to the healthy controls (Table 2), we analyzed the correlation between the miR-23a level and these parameters using Pearson Correlation Coefficient test. Data collected from the overall 6 groups (16 healthy controls and 56 CHDs patients) was used in the analysis. The results revealed that only CREA correlated with serum exosomal miR-23a level (r = 0.3255, *P* = 0.006) (Table 4 and Figure 5B), whereas HDLA and UA were not strongly associated with miR-23a level (Table 4, Figure 5A and Figure 5C).

**Table 4.** Correlation analysis between serum exosomal miR-23a levels and various clinical parameters of patients with CHDs. HDLC indicated High density lipoprotein cholesterol; CREA, serum creatinine; and UA, serum uric acid. \*\**P*<0.05

| Index | r value | *P* value |
| --- | --- | --- |
| HDLC | -0.09213 | 0.4481 |
| CREA | 0.3255 | 0.006** |
| UA | 0.03915 | 0.7531 |

**Figure 5.**
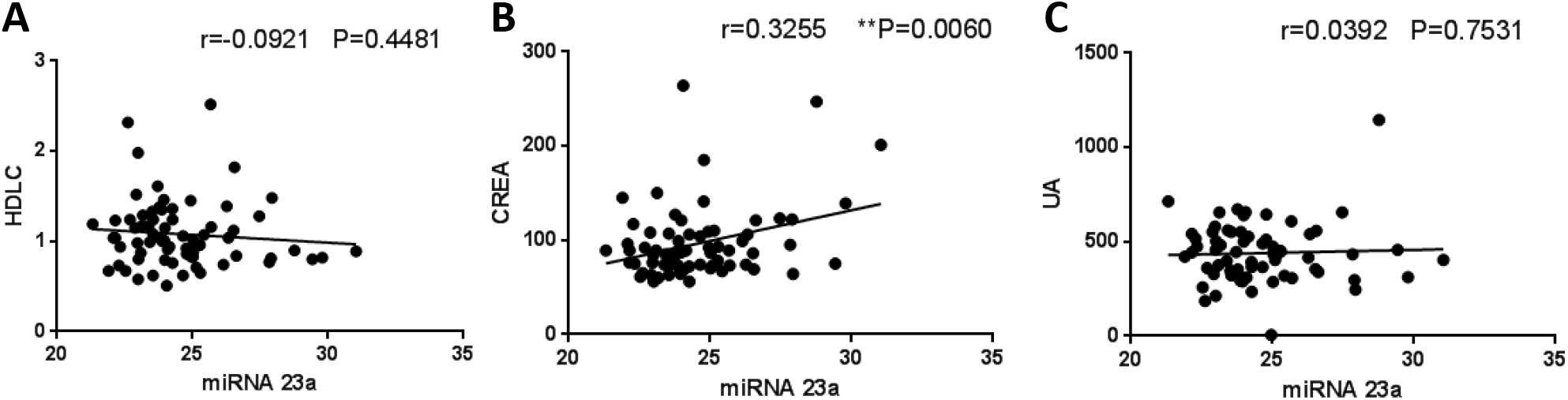
Correlation between serum exosomal miR-23a level and clinical parameters. **(A)** Scatter plot of the correlation between HDLC and miR-23a level. **(B)** Scatter plot of the correlation between CREA and miR-23a level. **(C)** Scatter plot of the correlation between UA and miR-23a level. \*\**P*<0.05

### Exosomal miR-23a was implicated in MI and CHDs

Considering the potential of miR-23a as a specific biomarker of stable angina and AMI, we further explored its roles in myocardial infarction (MI) and the overall CHDs. The MI group was a combination of AMI, STEMI and NSTEMI, including 32 MI patients in total. Compared with 16 healthy controls, the miR-23a level was also found downregulated in the MI population (Figure 6A). When we summarized all the CHD subjects, which contained the 56 patients from 5 CHD groups, the declined level of miR-23a was detected (Figure 6B). These results suggested that although miR-23a acted as a marker more specific to stable angina and AMI, and displayed no strong association with other CHDs in the current study, the significantly changed expression patterns in MI and the overall CHDs subjects suggested the miR-23a as a likely contributor to the disease mechanisms of CHDs.

**Figure 6.**
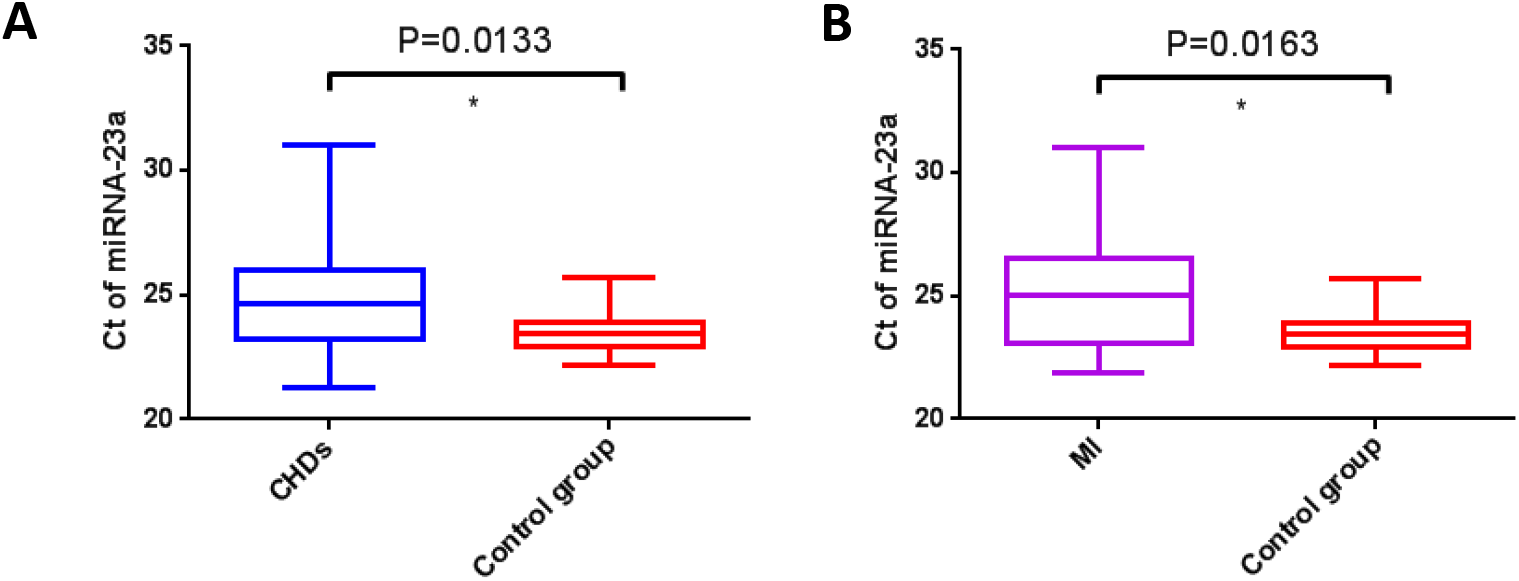
The expression of serum exosomal miR-23a was downregulated in the CHDs patients and in the MI patients. **(A)** Box plots of exosomal miR-23a expression in 16 healthy subjects and in 56 CHDs patients. **(B)** Box plots of exosomal miR-23a expression in 16 healthy subjects and in 32 MI patients. MI, myocardial infarction. \**P*<0.05

## Discussion

miR-23a/b had been implicated in various human diseases including cancers, cardiovascular diseases, muscle atrophy, sclerosis and metabolic diseases[23; 24]. The investigations of miR-23a were extremely focused on exploring its values as diagnostic and prognostic biomarkers for the carcinomas. So far, the miR-23a was demonstrated as overexpressed in the serum of cancer patients, including gastric, pancreatic, breast and colon cancer patients, and was strongly associated with specific clinical outcome of cancer patients[15]. Based on its expression pattern in various types of carcinoma, miR-23a alone or its combination with a panel of other miRNAs may serve as the non-invasive marker in cancer diagnosis[25]. Ogata *et al*. analyzed the exosomal miRNAs expression profile from serum of colorectal cancer patients and found the upregulated miR-23a in early stage patients, which proved that miR-23a was encapsulated in exosome indicating the potential of exosomal miR-23a in diagnosis[26].

Very limited studies were involved in assessing the expression pattern of miR-23a in cardiovascular diseases, meanwhile, the results seemed to be controversial. In the mouse model of apolipoprotein E deficiency (apoE−/−) induced coronary artery disease (CAD), miR-23a level was decreased in the mouse plasma[27]. And the downregulation of miR-23a was confirmed in the plasma from 32 CAD patients[27]. In another study, Satoh *et al*. analyzed the peripheral blood mononuclear cells (PBMC) taken from CAD patients and reported the overexpression of miR-23a in CAD-PBMC[28]. Our study revealed that miR-23a level was decreased in the serum exosome from 56 CHD patients (*P* = 0.0133) and 32 MI patients (*P* = 0.0163). In our further analysis of different characterized CHDs, miR-23a acted as a promising marker for the diagnosis of AMI and stable angina, whereas the miR-23a was not relative to STEMI, NSTEMI and unstable angina. Our data provided a novel understanding of the association of miR-23a and CHDs.

Similar as miR-23a, the abnormal expression of miR-23b was found in human hepatocellular carcinoma, lung cancer, colorectal cancer and ovarian cancer, and served as a promising diagnostic and prognosis biomarker[29; 30; 31; 32]. Exosome-derived miR-23b had been explored to be a novel therapeutic measure for preventing Kashin-Beck disease[33]. Blood miR-23b was found overexpressed in heart failure patients in a small-sample-size study of 8 healthy controls and 9 patients with congestive heart failure[34]. Interestingly, circulating miR-23b was reported as up-regulated in a comparison of 80 STEMI patients and 60 control subjects, and the increased miR-23b level declined in the blood samples of STEMI patients after percutaneous coronary intervention (PCI)[35]. We analyzed the serum exosomal miR-23b level of 8 control and 8 STEMI patients, and found the meanΔCt of STEMI group was lower than that of the control group (STEMI 23.6 vs. Control 24.2, *P* = 0.6294, Figure 2D), suggesting a likely trend of the up-regulation of miR-23b, whereas the change in miR-23b level displayed no statistical difference. Further study, however, would be required to amplify the sample size and demonstrate the association of miR-23b and CHDs.

miR-23a was demonstrated associated with CREA level of the overall CHD patients in the current study (*P* = 0.006). Meanwhile, the stable angina patients and AMI patients displayed more severe kidney injury with the significantly elevated CREA and UA levels and decreased miR-23a level, suggested that miR-23a might be involved in the CHD-renal damage. In the previous study, a diabetic animal model had proved that miR-23a combined with miR-27a attenuated the diabetes induced renal fibrosis, and the blood urea nitrogen (BUN) of diabetic animals was reduced after the miR-23a/27a intervention[23]. Interestingly, in a study of acute kidney injury (AKI) following AMI, the miRNA profile of AMI^+^AKI^−^ and AMI^+^AKI^+^ was compared, and the levels of miR-23a, miR-24 and miR-145 were shown significantly down-regulated in AMI^+^AKI^+^ patients compared to those in AMI^+^AKI^−^ patients[36]. Therefore the miR-23a has been revealed associated with kidney injury, and our data suggested a possible mechanism that miR-23a might mediate the CHDs induced renal dysfunction via the modulation of CREA-dependent signaling.

## Acknowledgements

This work was supported by the National Natural Science Foundation of China 81601317, the Natural Science Foundation of Guangdong 2017A030313584m, 2019A1515010290 (X. Yue).

## Conflicts of interest

None.

## Notes

### Competing Interest Statement

The authors have declared no competing interest.

